# Rotational dynamics versus sequence-like responses

**DOI:** 10.1101/2020.09.16.300046

**Authors:** Mikhail A Lebedev, Ivan Ninenko, Alexei Ossadtchi

**Affiliations:** Center for Bioelectric Interfaces of the Institute for Cognitive Neuroscience of the National Research University Higher School of Economics, Moscow, Russia; Department of Information and Internet Technologies of Digital Health Institute, I.M. Sechenov First Moscow State Medical University, Moscow, Russia

## Abstract

In a recent review, Vyas et al. commented on our previous observations regarding the presence of response sequences in the activity of cortical neuronal population and the contribution of such sequences to rotational dynamics patterns revealed with jPCA. Vyas et al. suggested that rotations generated from sequence-like responses are different from the ones arising from empirical neuronal patterns, which are highly heterogeneous across motor conditions in terms of response timing and shape. Here we extend our previous findings with new results showing that empirical population data contain plentiful neuronal responses whose shape and timing persist across arm-movement conditions. The more complex, heterogeneous responses can be also found; these response patterns also contain temporal sequences, which are evident from the analysis of cross-condition variance. Combined with simulation results, these observations show that both consistent and heterogeneous responses contribute to rotational patterns revealed with jPCA. We suggest that the users of jPCA should consider these two contributions when interpreting their results. Overall, we do not see any principal contradiction between the neural population dynamics framework and our results pertaining to sequence-like responses. Yet, questions remain regarding the conclusions that can be drawn from the analysis of low-dimensional representations of neuronal population data.

## Introduction

This study extends our previous report (Lebedev, Ossadtchi et al. 2019) where we conducted an analysis of multichannel recordings from monkey motor cortex, which Churchland, Shenoy and their colleagues had made available online (https://www.dropbox.com/sh/2q3m5fqfscwf95j/AAC3WV90hHdBgz0Np4RAKJpYa?dl=0) and which they had originally used to substantiate the concept of rotational dynamics – a fundamental type of neuronal processing in many species according to their claim (Churchland, Cunningham et al. 2012). In the rotational dynamics approach, multidimensional neuronal data are processed with a linear transform, called jPCA, designed to fit firing rates of the neurons in a population to their time derivatives and to generate a set of curved trajectories in a neuronal subspace, each trajectory corresponding to an arm-movement condition. (In their experiments, monkeys performed reaching movement with different paths.) If the rotational components obtained with this method account for a substantial percentage of variance in the data, this result is taken as a proof that the neuronal population acts as a dynamical system:

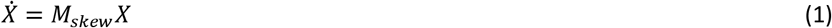

where *X* is a six-dimensional vector of principal components (PCs) obtained from neuronal population activity and 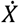 is its time derivative. The matrix *M*_*skew*_ has imaginary eigenvalues, which corresponds to *X* rotating around the center of coordinates (Garfinkel, Shevtsov et al. 2017).

**Figure 1.**
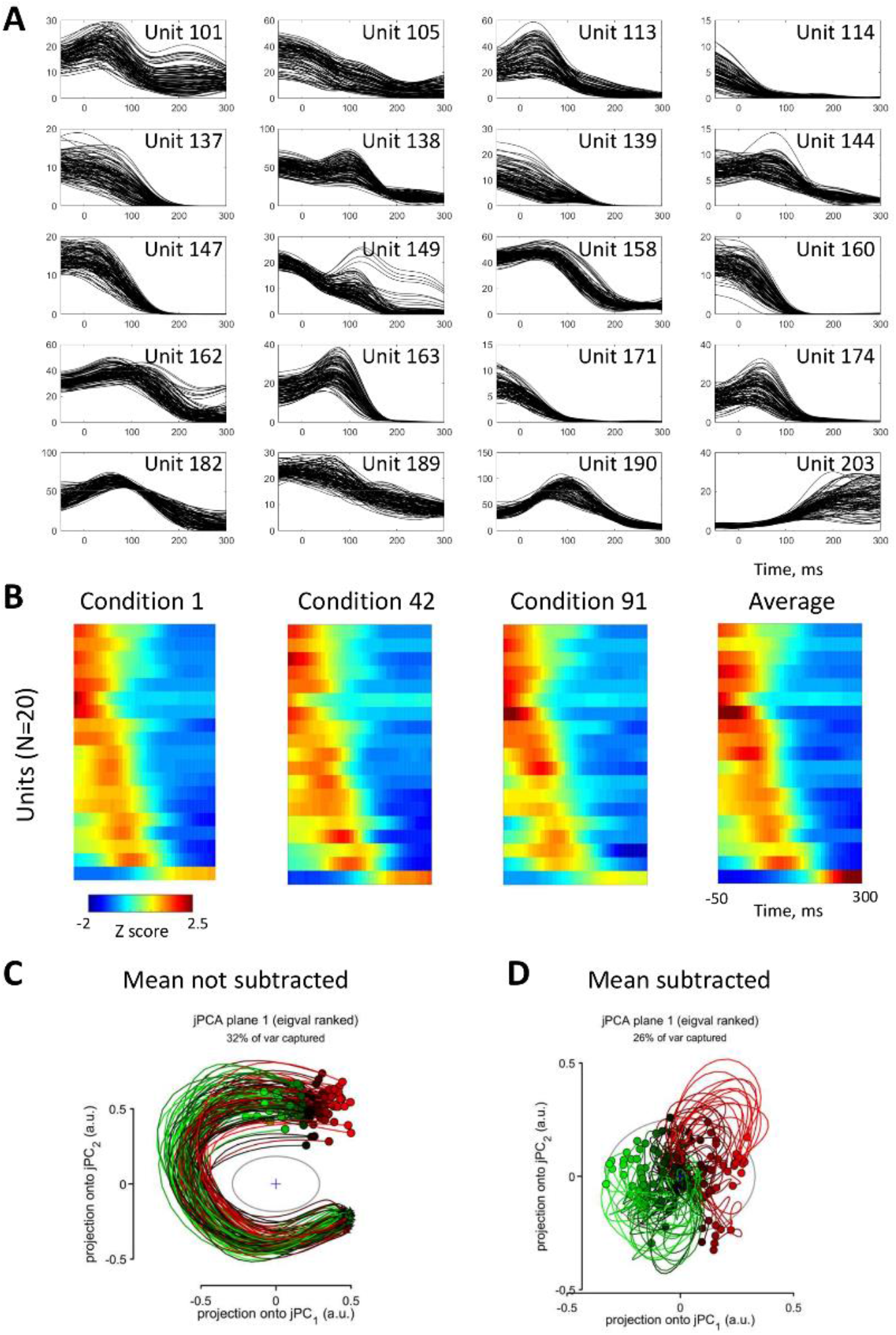
PETHs and jPCA plots for twenty neurons with the highest consistency of PETH shape across conditions. A: PETHs. Each box displays PETHs from different conditions for a single neuron. B: Population PETHs for three representative conditions (left) and the across-condition average PETH (right). Neurons are sorted by the time of peak response derived from the average PETH. C: State-space trajectories for jPCA without the subtraction of mean PETH. D: Trajectories for jPCA with mean PETH subtracted.

Given that *X*rotates, its coordinates are oscillating functions of time, and there is a temporal lag (or phase shift) between these functions. While Churchland and his colleagues obtained their results for PCs, we hypothesized that temporal lags could be present in the neuronal population data before it has been processed with jPCA (Lebedev, Ossadtchi et al. 2019). To test this hypothesis, we plotted population perievent time histograms (PETHs) for different conditions and sorted neurons by the time of peak activity estimated from the cross-condition average PETHs. With this simple procedure, we found a temporal sequence of neuronal responses during the arm reaching task. This sequence persisted for all 108 conditions with some variability (see Figure 2 in Lebedev, Ossadtchi et al. (2019)). Based on this observation, we argued that sequence-like responses could be the cause of the rotational dynamics revealed with jPCA. To tackle this idea further, we showed that rotational patterns could be easily obtained with jPCA from a rather arbitrary simulation of neuronal responses that incorporated a requirement that the responses be arranged in a temporal sequence but did not mimic any representation of movement. We concluded that jPCA reveals sequence-like patterns in neuronal population data, but we were skeptical regarding Churchland’s claim that this method elucidates a fundamental mechanisms of cortical motor control.

**Figure 2.**
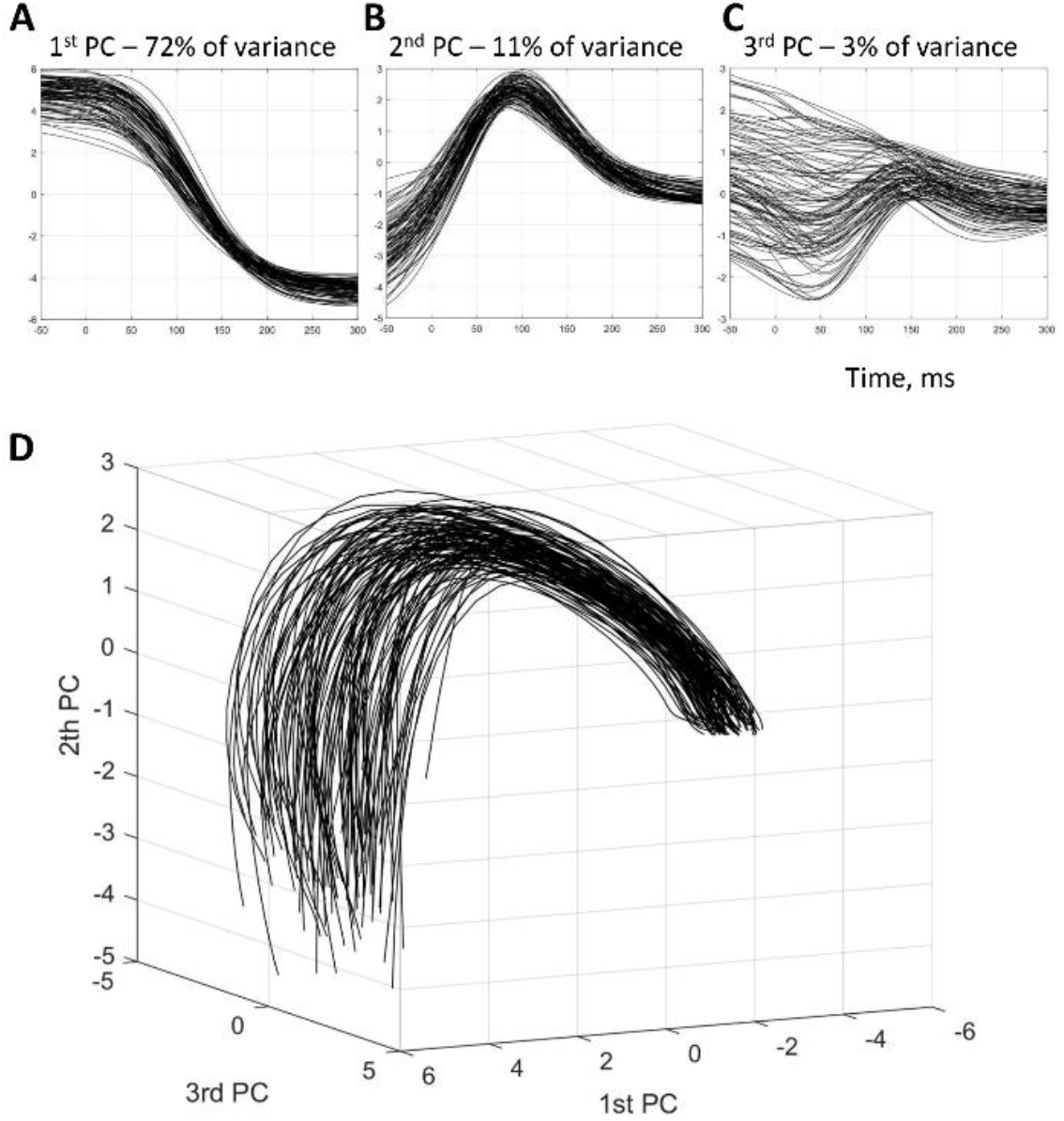
The first three principal components (PCs) for the population of twenty neurons with the highest consistency of PETH shape across conditions. A-B: The first, second, and third PCs. PETHs. D: The first three PCs in 3D.

Recently, the Shenoy group published a new review on neural population dynamics (Vyas, Golub et al. 2020) where they commented on our article:

> *“While it is possible to obtain rotation-like features from non-dynamical models [e.g., sequence-like responses (Lebedev et al. 2019)], such rotations are dissimilar from those observed empirically (Michaels et al. 2016). Perhaps more critically, the hypothesis of sequence-like responses does not account for single-neuron responses, which are typically multiphasic and/or peak at different times for different conditions (e.g., Churchland et al. 2012, figure 2).”*

These comments prompted us to provide several clarifications and conduct some additional analyses which we present here.

## Results

### Responses with consistent shape

We started our analyses with inspected the Shenoy group’s claim that “the hypothesis of sequence-like responses does not account for single-neuron responses, which are typically multiphasic and/or peak at different times for different conditions (e.g., Churchland et al. 2012, figure 2)”. Figure 2 in Churchland, Cunningham et al. (2012) indeed illustrates ten neurons (selected from the data of four monkeys) whose responses are highly variable across conditions in their shape and, consequently, peak timing. Yet, these types of responses could be considered as outliers since we have already shown that, at population level, a consistent sequence of neuronal responses is present for all conditions of Churchland’s data (see Figure 2 in Lebedev, Ossadtchi et al. (2019)).

To examine the issue of response shape consistency further, here we ranked the neurons in Churchland’s population by how regular PETH shape was across the conditions in different neurons. We quantified cross-condition consistency of PETH shape based on the correlation coefficients for PETH pairs representing different conditions for the same neuron. The mean of these correlation coefficients was used as a measure of PETH shape consistency for each neuron. This metric detected persistent PETH shapes while ignoring the differences in amplitude. The mean correlation coefficient ranged from 0.005 to 0.988 for different neurons (N=218), with the median of 0.494.

The neurons with high cross-condition consistency of PETH shape fit the pattern of “sequence-like responses” the best because in such neurons peak activity occurs at a constant time with respect to movement onset. Provided there is a spread in the occurrences of peak responses in different neurons, the peaks form a sequence. To assess jPCA results for the neurons best fitting this description, we selected top twenty neurons with the highest mean correlation coefficients for PETH pairs (Figure 1). Mean correlation coefficient was greater than 0.90 for all the neurons selected. Figure 1A shows these neurons’ PETHs; traces for different conditions are superimposed. Clearly, PETH shapes were highly consistent across the conditions for these neurons while there was some cross-condition variability in response amplitude. Furthermore, after we sorted these neurons by the time of peak response, a temporal sequence of activity was clear, which was preserved across the conditions (Figure 1B).

The originally proposed jPCA method includes two options: the option where cross-condition mean PETH is subtracted from each PETH before the computation of PCs and the one where no such subtraction is performed. For the twenty neurons with the most consistent PETH shapes, we ran jPCA with the subtraction (Figure 1D) and without it (Figure 1C). In both cases, jPCA yielded curved trajectories. The rotations were especially clear and similar for all trials when mean PETH was not subtracted, whereas at least two clusters of curved trajectories (red and green traces generated by Churchland’s script) were produced by jPCA when mean PETH was subtracted. Since the subtraction of mean PETH distorts response shape (Lebedev, Ossadtchi et al. 2019), it is not clear whether these two clusters represent any meaningful result.

We next ran a principal component analysis (PCA) for the same twenty neurons (Figure 2). The responses were highly consistent across the conditions when expressed as the first (Figure 2A) and second (Figure 2B) principal components (PCs), which explained 72% and 11% of the data variance, respectively. The responses for the third PC (Figure 1C), which corresponded to 3% of the variance, were variable across the conditions. The shapes of the first, second, and third PCs were approximately sinusoidal and were shifted in time against each other, which resulted in the curved trajectories in the 3D plot with these PCs at the axes (Figure 2D). The PC trajectories highly resembled the traces obtained using jPCA without the subtraction of mean PETH (Figure 1C). Similar results were obtained when more than twenty neurons with consistent PETH shapes were selected for this analysis (e.g., 50 or 100) or when neurons ranked 21 through 40 were selected, 41 through 60, and so on until 81 through 100. When the neurons with less consistent PETH shapes were used, the first two PCs became more variable across the conditions and the sequence of peak responses visible in the population color plots became noisier, and, approximately after the middle of the population was crossed, the sequence vanished.

Thus, Churchland’s data contained a sizeable subpopulation of neurons with consistent PETH shapes and, consequently, consistent sequence-like responses, so the statement that single-neuron responses are “typically multiphasic and/or peak at different times for different conditions” is inexact.

### Heterogeneous responses

The neurons with heterogeneous responses, like the ones illustrated in Figure 2 of Churchland, Cunningham et al. (2012), could be found in Churchland’s data using the same criterion of PETH shape consistency as we used in the previous analysis. To examine the neurons of this type, we selected 20 neurons with the least consistent PETH shapes (Figure 3). The variability of PETH shapes can be appreciated from Figure 3A. Although the responses were variable across the conditions, they were not completely void of a temporal sequence. For example, in Figure 3A, firing rate of unit 3 was modulated earlier compared to unit 165, and epochs of high rate variability can be noticed for the other units, as well. For the response patterns of this kind, the sequence of responses could be characterized by calculating cross-condition PETH variance as a function of time for each neuron, sorting the neurons by the time of peak PETH variance, and then displaying the time-dependent PETH variance as a colorplot (Figure 3B). To be precise, for better visibility, Figure 3B shows the standard deviation, i.e. the square root of variance. In this plot, peaks correspond to high variability of neuronal activity across the arm-movement conditions. High movement-dependent variability is not surprising for motor cortical activity, although the fact that the timing of this variability is fairly consistent in individual neurons can be considered a novel finding.

**Figure 3.**
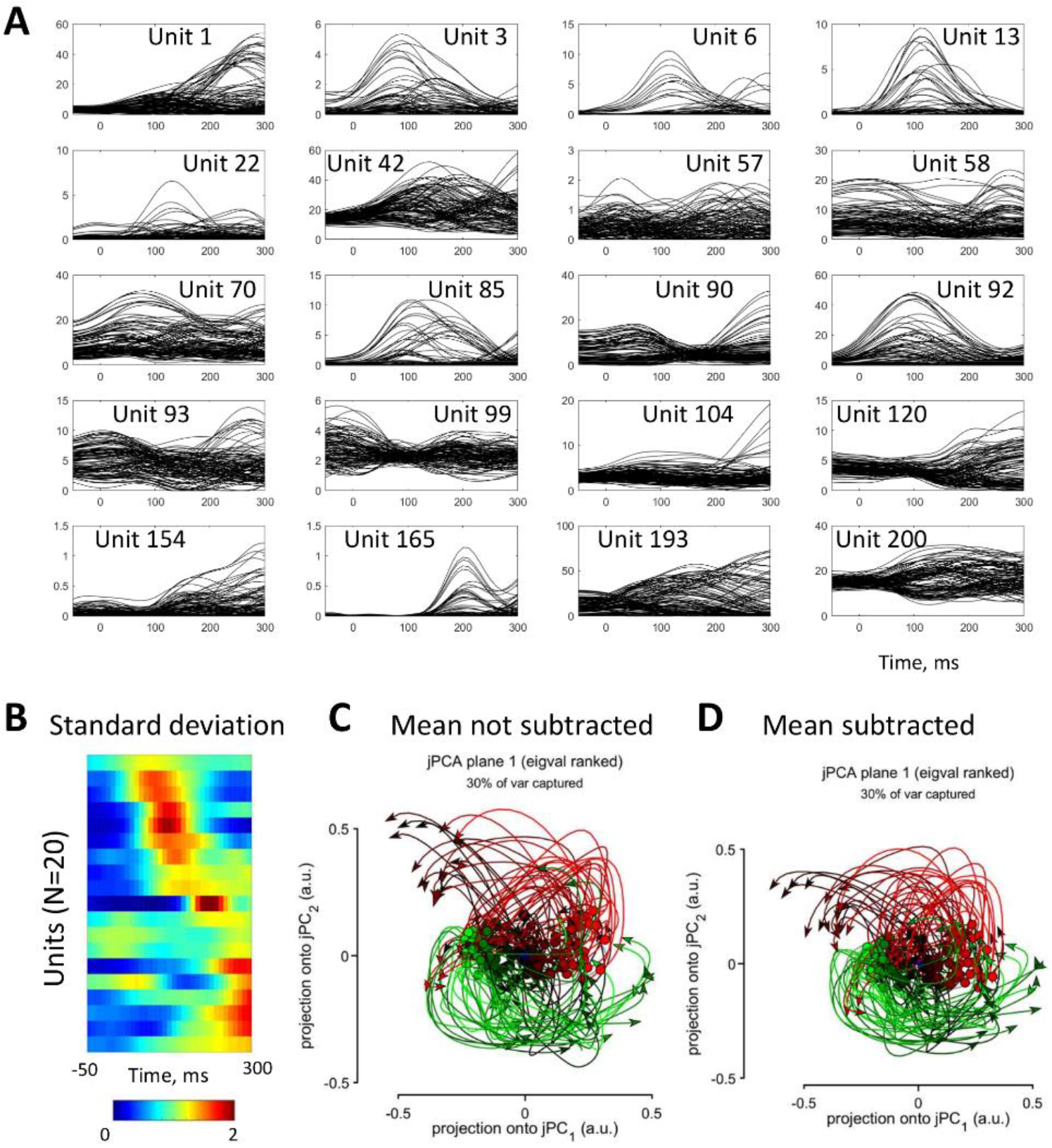
PETHs and jPCA plots for twenty neurons with the lowest consistency of PETH shape across conditions. A: PETHs. Each box displays PETHs from different conditions for a single neuron. B: Color plot of cross-condition PETH variance. Neurons are sorted by the time of peak variance. C: State-space trajectories for jPCA without the subtraction of mean PETH. D: Trajectories for jPCA with mean PETH subtracted.

The jPCA conducted for the subpopulation of twenty neurons with highly variable responses yielded two clusters of curved trajectories (red and green lines in Figure 3B,C) that were only moderately affected by the subtraction of mean PETH. The PCA conducted for the same neurons (Figure 4) revealed high cross-condition variability for each of the first three PCs (Figure 4A-C), which accounted for 17, 11, an 9% of the variance in the data. Clusters of conditions with similar responses could be noticed in these plots, as well as in the 3D plot for the first three PCs (Figure 4D). The phase shifts between the PCs accounted for the curvature in the plots.

**Figure 4.**
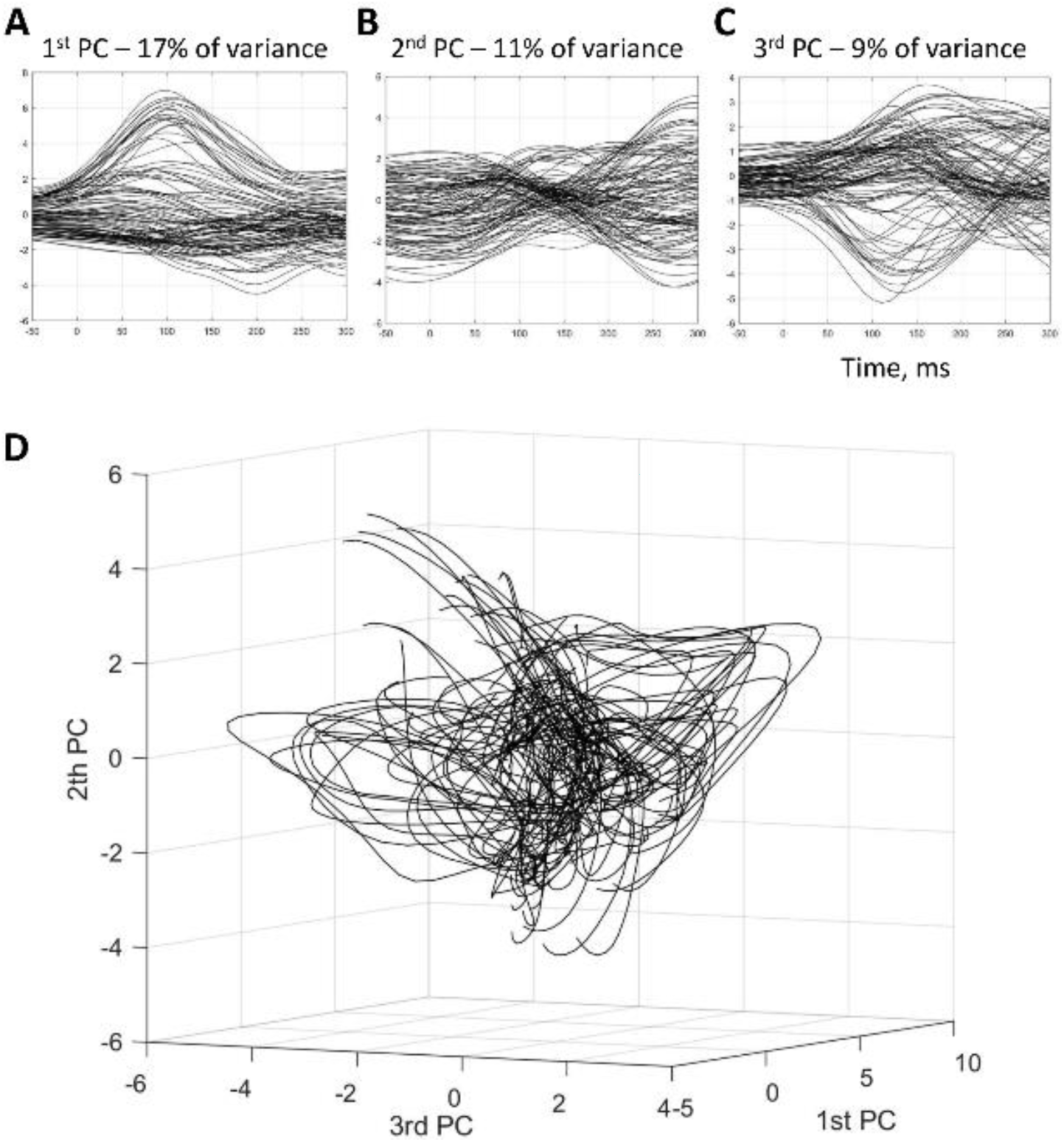
The first three principal components (PCs) for the population of twenty neurons with the lowest consistency of PETH shape across conditions. A-B: The first, second, and third PCs. PETHs. D: The first three PCs in 3D.

Thus, while neurons with heterogeneous response were present in Churchland’s neuronal population, these neuronal modulations were non-uniformly distributed in time, with some neurons being modulated earlier than the others. Notwithstanding the high variability of responses in individual neurons, these responses were sufficiently correlated across the neurons, as evident from the contribution of the first three PCs to the variance (37%; compare to 86% for the neurons with consistent PETH shapes), which assured that jPCA analysis reliably captured the population responses. Then, because the sequence of responses phase shifts between the PCs originated from the sequence of PETH variance and eventually resulted in the curved jPCA trajectories.

### PETH peaks versus variance

Since, in addition to the definition of response sequence based on the occurrences of PETH peaks (Figiure 1B; also see Figure 2 in Lebedev, Ossadtchi et al. (2019)), we introduced the definition based on the timing of cross-condition variance (Figure 3B), it was of interest to compare these two definitions. We used both approaches for the entire population of Churchland’s neurons and found similarities in the resulting sequences (Figure 5). In Figure 5A, the sequence was derived from PETH variance (shown on the left) and used for sorting neurons in the population PETH plot (shown on the right). The sequence of PETH peaks was still present after this operation. Conversely, defining the sequence based on the timing of PETH peaks (Figure 5B, right) preserved the sequence in the plot showing cross-condition variance (Figure 5B, left). Thus, the average response was roughly proportional to average variance.

**Figure 5.**
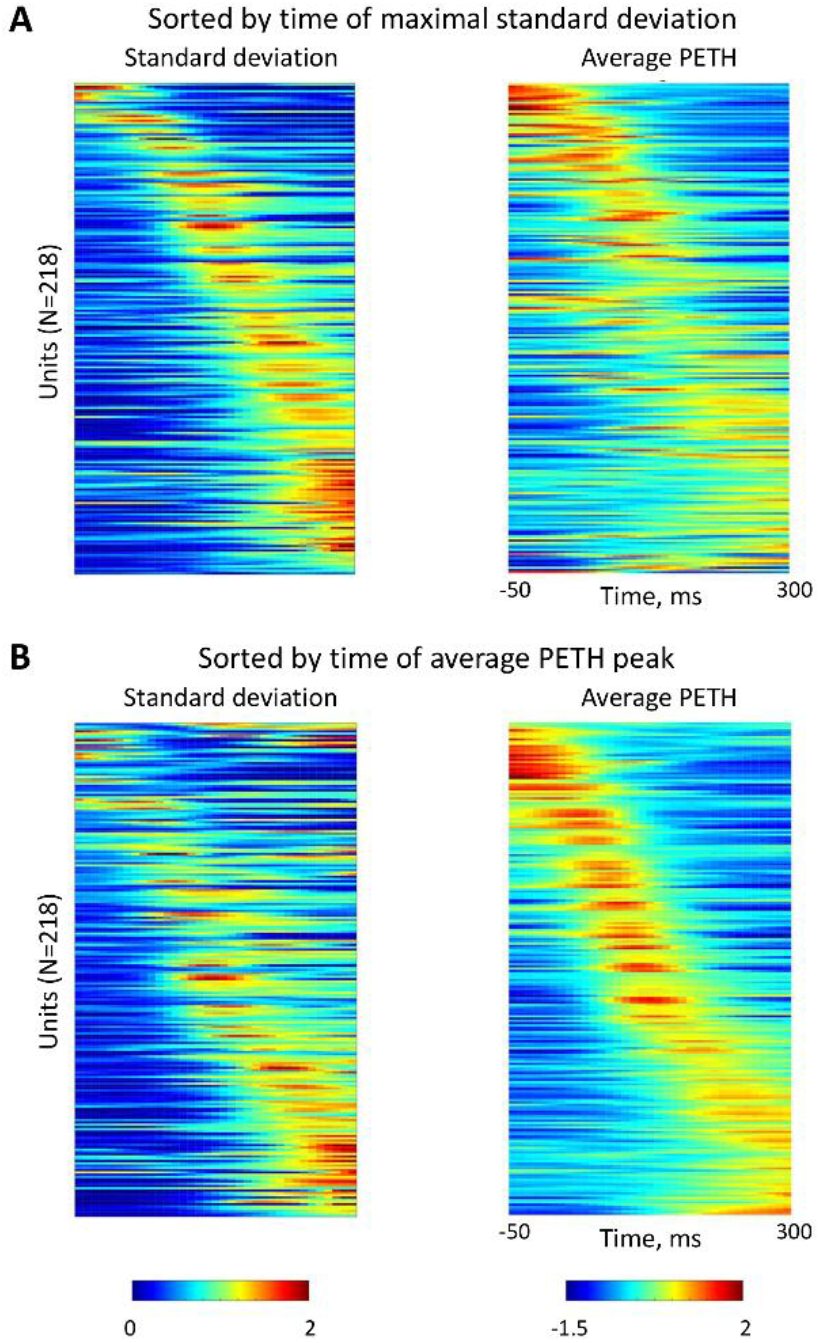
Response sequences defined by the time of maximal PETH variance or PETH peak. A: The sequence was derived from PETH variance (left) and then applied to average PETHs (right). B: The sequence was derived from average PETHs (right) and then applied to PETH variance (left).

### Simulations

To verify the mechanism that we proposed for explaining the emergence of rotational patterns in jPCA plots for the neurons with heterogeneous responses, we ran a simulation (Figure 6), where conditions (N=108) represented directions of center-out movements (from 0 to 360 degrees) and neurons (N=218) were modeled as having cosine tuned directional tuning. A similar simulation was previously reported by Michaels and his colleagues (Michaels, Dann et al. 2016), although they did not discuss response sequences explicitly and did not analyze the effect of removing the sequence while keeping the other response parameters the same. In our simulation, the same shape of PETH was chosen for all neurons (Figure 6A), and its amplitude was cosine-tuned to movement angle. Preferred directions were randomly selected from the range 0 to 360 degrees (Figure 6B). We first simulated a population with responses in individual neurons occurring simultaneously (Figure 6C). For this population, rotations were found neither in the plot depicting three initial PCs (Figure 6D) nor the jPCA plot (Figure 6E).

**Figure 6.**
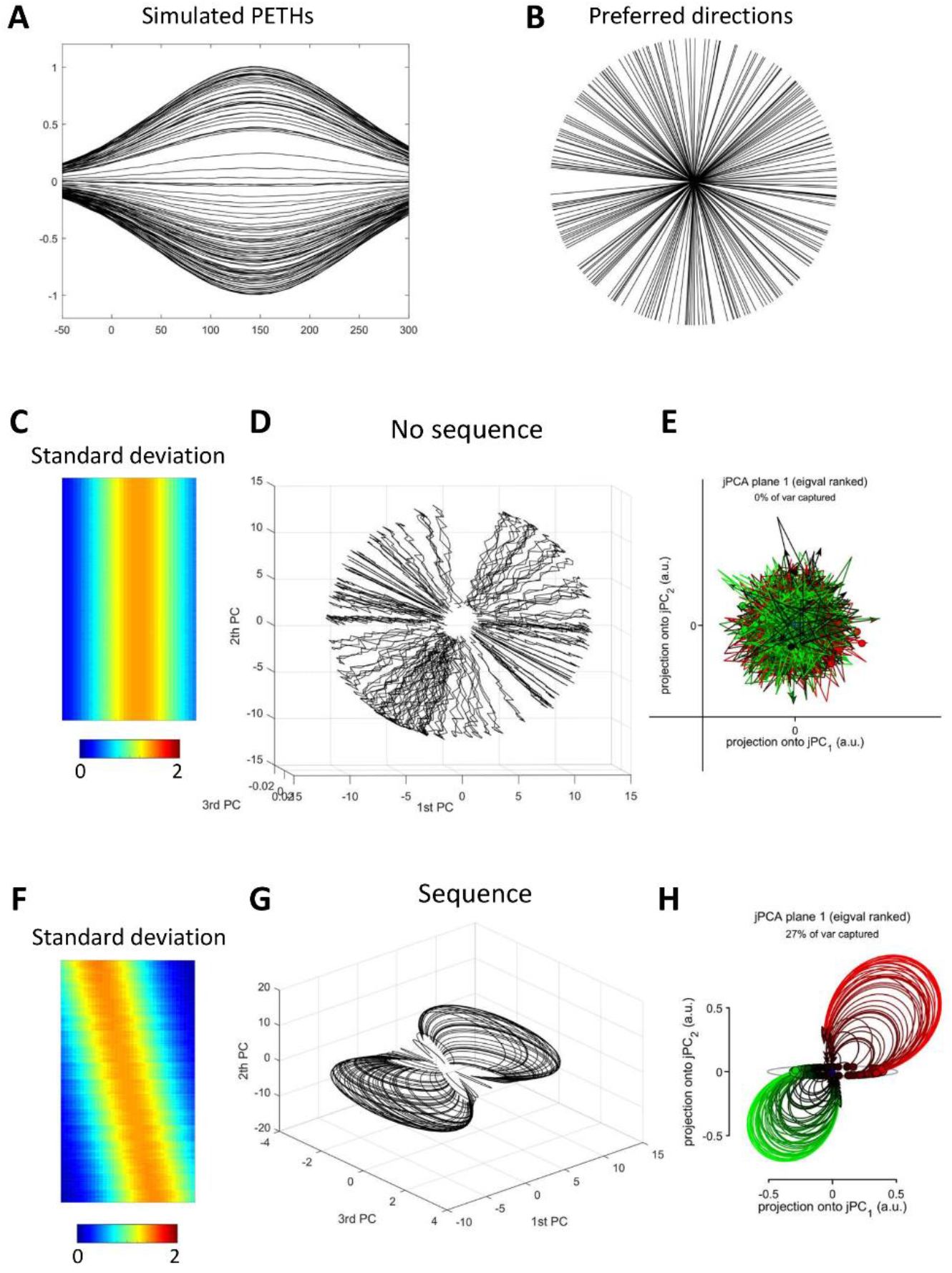
Simulations of cosine-tuned neurons with and without response sequence. A: PETHs for one neuron for different conditions (movement directions). PETH amplitude is proportional to movement angle. B: Preferred directions for the neurons in the population. C: Cross-condition standard deviation. In this population there was no response sequence. D: 3D plot of the initial three PCs. E: jPCA plot. F-H: the same analyses for the simulation of a response sequence.

Next, we simulated a neuronal population with a sequence of responses (Figure 6F). The preferred directions were kept the same as in the previous analysis. For this population, rotations were visible in both the plots depicting PCs (Figure 6G) and the jPCA plot (Figure 6H).

## Discussion

Here we conducted additional analyses in response to the comments of Vyas, Golub et al. (2020) regarding our paper (Lebedev, Ossadtchi et al. 2019). Although Vyas et al. asserted that we proposed sequence-like responses as an alternative to dynamical models of population activity, this is not what we suggested. We did not advocate any non-dynamical model that would oppose the dynamical-system framework of Shenoy and his colleagues. As a matter of fact, we did not propose any physiologically plausible neuronal model but instead analyzed the empirical data of Churchland and his colleagues or ran simulations that were not based on any physiological findings. We started with analyzed Churchland’s data, where we found evidence of sequence-like responses. This finding was not new; several previous studies reported similar neuronal patterns (Georgopoulos, Kalaska et al. 1982, Luczak, Barthó et al. 2007, Lebedev, O’Doherty et al. 2008). The novel finding of ours was the observation of a response sequences that was preserved across different movement conditions. While it was tempting to interpret such a sequence in terms of consecutive stages of neuronal processing, we thought that the dynamical-system explanation proposed by the Shenoy group was plausible, as well.

As to the comparison of representational and dynamical models conducted by Michaels, Dann et al. (2016), we consider their work an effort in the right direction but we have little to add to this topic simply because we did not examine how the neurons in Churchland’s dataset encode motor parameters. (And we could not because the dataset does not contain kinematics data.) Yet, there appears an important point of agreement between us and Michaels and his colleagues, as they concluded that “incorporating variable latency offsets between neural activity and kinematics is sufficient to generate rotational dynamics at the level of neural populations”. The same judgment was expressed by the reviewer of our manuscript who wrote that it was obvious that “state-space rotation is equivalent to a preserved sequence of neural activity”. Since there appears to be a consensus on the issues of sequences producing rotations and sequences being present in empirical data, we suggest that “sequence-like responses” should not be considered as something completely different from a “dynamical model” or an “empirical observation”. We instead suggest that an investigation of response sequences should be viewed as a complementary analysis to the population methods that reduce data dimensionality, such as jPCA.

Looking into how response sequences (taken from experimental data or simulated) could be helpful to validate the results obtained with jPCA and similar methods, which are often hard to interpret. For instance, we described cases where there was a clear sequence in neuronal responses but jPCA did not return rotations because it incorporated a subtraction of cross-condition mean response (Lebedev, Ossadtchi et al. 2019). Algorithmic components of this kind could significantly alter the results an mislead a researcher that relies exclusively on a complicated population analysis. For instance, Michaels, Dann et al. (2016) permutated their data and then generated rotational patterns – a procedure that could have resulted data that was ill conditioned for jPCA. Yet, they drew their conclusion from the state-space plots.

The analyses that we conducted here provide additional explanations to how sequence-like responses contribute to what Churchland and his colleagues called rotational dynamics (Churchland, Cunningham et al. 2012). They used jPCA method to extract rotational patterns from neuronal population activity and argued that the existence of such patterns constitutes a proof that monkey motor cortex acts like a dynamical system. While the dynamical system idea is currently popular, we think that the results generated with dimensionality reduction methods need scrutiny, particularly with respect to their relationship to the activity patterns of individual neurons within the population.

In their response to our work, the Shenoy group was skeptical regarding response sequences being the main cause of rotational dynamics. They also called sequence-like responses a non-dynamical model even though their own data contain such sequences. In this paper, we provided additional evidence showing that Churchland’s dataset contains neurons whose response timing is highly consistent across conditions (Figure 1). Previously the Shenoy group emphasized that motor cortical neurons have heterogeneous response patterns and did not describe the neurons with consistent responses.

For the highly consistent neurons, a sequence of responses can be easily revealed with a simple analysis of response peak timing. These neurons fit Churchland’s original description of “an unexpected yet surprisingly simple structure in the population response” as they exhibit remarkably similar responses for all arm-movement conditions. Yet, Churchland, Shenoy and his colleagues have expressed dissatisfaction with this simple population pattern (Churchland, Cunningham et al. 2012) because it corresponds to “rotations that are similar for all conditions”. They reasoned that “in such cases one fails to gain multiple views of the underlying process, making it difficult to infer whether rotations are due to dynamics or to more trivial possibilities”. We respectfully disagree with this assessment because, in our opinion, cross-condition consistency fits the dynamical system framework quite well. For example, Figure 1C depicts state-space trajectories for the neurons with consistent responses. These trajectories resemble the state diagram for a pendulum (see Figure 1c in (Vyas, Golub et al. 2020)), and even a dependence of the trajectory on the initial state can be noticed. Curiously, these neurons resemble the neurons that Michaels and his colleagues produced with their dynamical model (Michaels, Dann et al. 2016).

The other point of disagreement is related to the way Churchland et al. dealt with “rotations that are similar for all conditions”. To remove these rotations, they subtracted cross-condition mean PETHs from the PETHs for each condition. In the case of the neurons with consistent responses, this procedure simply removes most of the data and leaves noise (e.g. compare Figure 1C and Figure 1D), so it is unclear what one is supposed to gain with this method. The gain is not clear either for the neurons that are “multiphasic and/or peak at different times for different conditions”. In this case, mean PETH is likely to be highly noisy. Additionally, mean PETH could be dominated by the strongest responses contributed by a few conditions (e.g. preferred-direction response). In addition to these considerations, it is unclear how a dynamical system defined by equation 1 could perform subtraction of the mean PETH.

The point the Shenoy group made about the neurons with heterogeneous responses merited a special consideration. A sequence of responses was not clear for the neurons of this type was not clear when looking at the sequence of PETH peaks. Yet, after we defined the sequence using the timing of PETH variance, it became clear that neuronal modulations did not occur randomly in time but followed a sequence instead. This sequence eventually translated into the curved jPCA trajectories.

Overall, we do not see any principal controversy between the neural population dynamics framework (Vyas, Golub et al. 2020) and our results pertaining to sequence-like responses. Yet, in our opinion, questions remain regarding the interpretation of low-dimensional representations of neuronal population data.

## Methods

The data and jPCA code were downloaded from the Churchland lab’s online depository

(https://www.dropbox.com/sh/2q3m5fqfscwf95j/AAC3WV90hHdBgz0Np4RAKJpYa?dl=0). The dataset contained PETHs for each neuron and each condition. jPCA was run with the following MATAB commands:

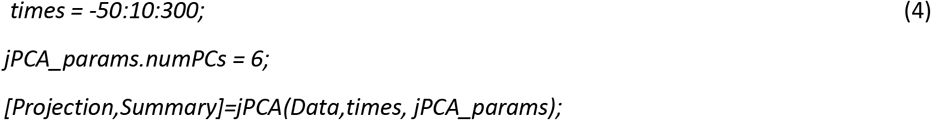

PETHs were standardized by subtracting the mean (for all conditions) and dividing by the standard deviation (also for all conditions).

## Acknowledgement

This work was supported by the Center for Bioelectric Interfaces of the Institute for Cognitive Neuroscience of the National Research University Higher School of Economics, RF Government grant, ag. No. 14.641.31.0003.

